# Lexical surprisal drives pupil dilation during continuous story listening

**DOI:** 10.64898/2026.10.26.761911

**Authors:** Chiara Terzo, Floris P. de Lange, Eva Berlot

## Abstract

The pupil famously adjusts its dilation in response to light, but its role extends far beyond this basic sensory function. Its fluctuations index arousal and mental effort, beyond luminance-related dynamics, making it a sensitive marker of internal cognitive states. A growing body of work shows that these pupil-linked signals extend beyond the visual domain. However, which processes drive pupil responses in more naturalistic settings remains unclear as most evidence comes from tightly controlled paradigms that strip away the richness of real-world experience. Language offers a uniquely powerful window into this question, as it can allow us to create rich paradigms without having any visual, i.e. luminance, confounds. Moreover, language comprehension hinges on continuous prediction: listeners constantly generate expectations about upcoming input, and the discrepancy between expectation and actual input - contextual prediction error, or surprisal - carries measurable cognitive costs. Here, we tested the hypothesis that pupil-linked arousal indexes these linguistic surprisal signals during naturalistic listening. We recorded pupil responses while volunteers listened to naturalistic spoken stories in darkness, which were either intact or reversed. Reversed speech preserves the acoustics of the signal while destroying its intelligibility. Pupil dilation selectively tracked lexical surprisal within a late window of 1,700 to 2,350 milliseconds following word onset, in forward, but not control, backward speech. These findings indicate that pupil-linked arousal is tuned to global, context-dependent processing, tracking the cognitive cost of updating a predictive model during naturalistic listening.

## INTRODUCTION

The pupil is classically understood as a regulator of light entering through the retina. Through the pupillary light reflex, pupil diameter constricts under increased luminance thereby controlling the amount of light reaching the retina^[1,2]^. The pupillary light reflex is not, however, exclusively driven by physical luminance. High-level visual interpretation can modulate it. For example, viewing photographs of the sun elicits pupillary constriction relative to luminance-matched control images, indicating that semantic content can influence the pupil^[3]^.

Beyond these luminance-related dynamics, the pupil has emerged as a sensitive marker of internal cognitive states within the visual domain. It tracks covert attention^[4,5]^, mental imagery^[6,7]^, implicit learning of visual regularities^[8,9]^, and novelty during visual encoding^[10]^. Task-evoked pupillary dilation in these contexts exhibits two temporally and functionally distinct signatures: a rapid, transient phasic response to unexpected or salient events, and a sustained, graded dilation that scales proportionally with cognitive load and emotional engagement^[11-15]^. The former reflects processes of novelty detection and surprise, putatively priming the system for subsequent exploration; the latter, mediated by parasympathetic inhibition, intensifies under conditions of escalating mental effort^[16]^.

A critical limitation, however, is that these effects have been documented almost exclusively in visual paradigms. In such settings, pupillary responses remain vulnerable to confounds from low-level changes in luminance and contrast. Even when stimuli are carefully matched for mean luminance, local contrast fluctuations can drive pupil changes independently of the cognitive or predictive processes under investigation. This motivates the study of pupil dynamics in non-visual modalities, where retinal illumination can be held constant and cognitive effects isolated.

Auditory research circumvents this issue, allowing for cleaner isolation of prediction-related dynamics. In purely auditory paradigms, pupil dilation is consistently driven by deviations from an established acoustic regularity^[17]^, and is sensitive to multiple hierarchical levels of deviation, distinguishing low-level sensory surprisal from higher-order contextual shifts^[18]^. Likewise, in an auditory predictive-inference task, Filipowicz et al.^[19]^ found that baseline pupil size encoded belief strength while evoked responses tracked surprise after prediction errors. These dynamics varied with individual differences in belief updating, especially when predictions and sensory evidence diverged, suggesting that pupil arousal indexes covert exploration toward uncertainty reduction, bridging mental effort and computational inference. Across both visual and auditory modalities, these findings converge on the idea that pupil size tracks surprise and uncertainty, scaling with the degree of expectation violation. Within the predictive coding framework^[20,21]^, such responses are understood as markers of elevated arousal that prompt the brain to revise its ongoing predictions ^[9,17,18,22-28]^.

Building on this, we next consider whether pupil dynamics scale with the predictive demands of more naturalistic stimuli. Spoken language is a particularly well-suited test case: it is hierarchically organized, unfolds in real time, and places sustained predictive demands on the listener, forcing the transformation of an ambiguous acoustic stream into increasingly abstract representations of meaning^[29-31]^. Indeed, pupil diameter has already been used as a metric of listening effort in naturalistic contexts such as continuous narratives^[32]^, and recent work shows that during narrative listening it tracks surprisal-based prediction signals when analyzed with temporal response functions^[33]^. Accordingly, linguistic processing demands can be conceptualized along a continuum from local to global levels of representation. At the most global level, the brain generates expectations about upcoming input based on extended linguistic context. Two complementary metrics capture distinct aspects of this process: surprisal, which quantifies the local unexpectedness of an observed word given its preceding context, and entropy, which reflects the uncertainty of the context before a word is encountered^[34-36]^. Surprisal measures the actual prediction error for a specific word, whereas entropy measures the expected uncertainty of the preceding context; high entropy indicates an unpredictable context, low entropy a highly predictable one. Neurophysiological evidence indicates that the brain distinguishes between static semantic fit and dynamic contextual predictability across hierarchically organized levels of linguistic representation^[30,37,38]^, suggesting that these different types of information engage distinct neural mechanisms.

As lower-level controls, two further metrics index processing demands that are independent of this dynamic predictive machinery. At a broader level of representation, semantic integration, the process of combining an incoming word’s meaning with preceding discourse to form coherent mental representations^[39]^, reflects the degree to which a word fits within the overall semantic fabric of its preceding context, operationalizable as static, context-independent semantic similarity. At the most local level, lexical access itself imposes processing demands: word frequency, a direct index of lexical difficulty, drives pupil dilation even in single-word paradigms where contextual integration is minimal^[40,41]^, reflecting the retrieval of a word’s form and meaning from long-term memory.

This raises an open question: is the pupillary system sensitive to the entire hierarchy of linguistic processing demands during continuous narrative listening, spanning local lexical access and semantic integration through global contextual prediction? To address this, we employed an auditory narrative listening paradigm in which participants listened to podcasts while their eye movements and pupillometric data were recorded. We used large language models to quantify how linguistic surprisal and predictability changed with the narrative, and investigated to what extent the pupil tracked these distinct levels. As a control condition, we also played the same podcasts in reverse: this preserves the acoustic envelope and other surface properties of the speech signal, while fully disrupting narrative/language understanding. To preface the results, we found that the pupil selectively tracked lexical surprisal, specifically in forward speech.

## METHODS

### Participants

A total of 49 participants took part in the study. One participant withdrew at the start of the experimental session due to adverse events, while data from five participants could not be utilized because of technical issues and poor eye-tracking signal, resulting in a final sample of 43 participants (10 males, 33 females, mean age=21.81, range 18-29 years). All participants were native Dutch speakers, had normal vision and provided informed consent prior to their involvement in the study. All procedures were approved by the local ethics committee [Commissie Mensgebonden Onderzoek (CMO) regio Arnhem-Nijmegen, The Netherlands].

### Stimuli and procedure

Participants listened to continuous narrative speech from two Dutch audio stories taken from the podcast Echt Gebeurd (https://echtgebeurd.net/), each lasting approximately 8 minutes. Both stories were presented twice. As a control condition, one of the stories was played backwards, also presented twice. The backward condition served as a control in which the low-level acoustic properties (e.g. acoustic envelope) are preserved, just reversed, while the high-level intelligibility of the content is fully disrupted (Figure 1). The order of the stories and the choice of podcast for backward story were all counterbalanced across participants. In total participants listened to six runs: Story 1 and its repetition, Story 2 and its repetition, and the backward version of one of the stories and its repetition. All three stories were first presented once before they were all repeated. All stimuli were presented through two speakers, placed equidistantly from the participants on the left and right sides of the table, with a sampling rate of 44.1 kHz.

**Figure 1.**
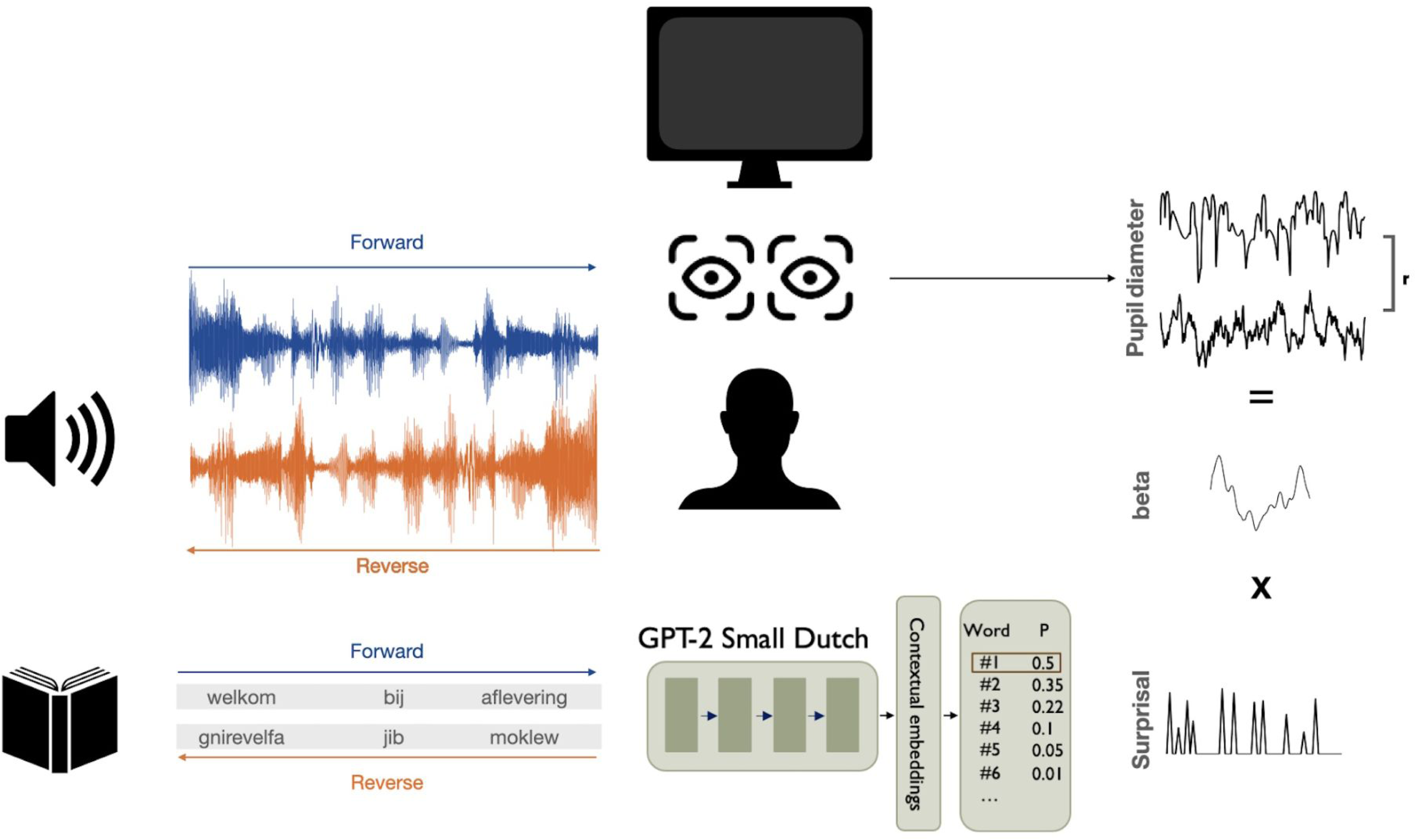
Experimental paradigm and analysis. Participants listened to two audiostories taken from the Dutch podcast Echt Gebeurd while their eye movements and pupil dilation were recorded. The audiostories were played twice each, and one story was also played temporally reversed, twice. A Dutch GPT-2 model was applied to the audiobook transcript to quantify contextual surprisal for each word. Linear relationships between linguistic features and the pupil signal were estimated using temporal response function (TRF) modeling. The TRF model computes the beta weights for each time lag, isolating the distinct contribution of surprisal feature to the pupil signal. Performance was assessed using cross-validation, with prediction accuracy expressed as the Pearson correlation (r) between model-predicted and empirically observed pupil signals on held-out data.

The experiment was conducted in a dark room, with participants seated in front of a black screen set to the lowest luminance. To enhance the darkness of the environment and minimize any luminance effects on pupil response, a black board was placed behind the screen. This arrangement ensured that participants were not distracted by any objects in their line of sight, allowing for natural and unrestricted eye movements. At the beginning of each run, participants were exposed to a brief 15-second baseline resting period, during which they were instructed to sit quietly. The only instruction given was to listen attentively to the podcast. While participants were encouraged to move their eyes naturally, they were advised to avoid abrupt movements of their body or head to ensure accurate eye tracking. At the end of each run, participants listened to two 8-second audio segments from the same recording they had just listened to, separated by a 4-second interval. They were instructed to press the left button on a response pad if they believed the second segment was a continuation of the first, or the right button if it was not. This prompt served to additionally motivate participants to listen attentively.

### Audio preprocessing

The audio files were preprocessed using the PRAAT software package^[42]^ to trim unnecessary sections from the beginning and end of each experimental run/block. The reversed audio version was generated using custom MATLAB (R2021b MathWorks) code.

### Stimulus annotations

Audio files and their transcripts were aligned via the Montreal Forced Aligner^[43]^. Forced alignment is a technique that takes an audiofile’s orthographic transcription and produces a time-aligned version by using a pronunciation dictionary to map words to their corresponding phonetic units. The aligner, pre-trained on a Dutch dictionary, processes both the audio file and the transcript’s text grid to generate an alignment. This alignment was then imported into PRAAT for manual review, focusing on correcting any misalignments caused by words not included in the aligner’s dictionary. Finally, word onset and offset timestamps were extracted through PRAAT for each story. For backward speech, ‘word onset’ was determined by reversing forward-to-backward the word offsets from the forward stories.

### Eye movements acquisition and pre-processing

#### Acquisition

Eye movements and pupil data from both eyes were recorded using an EyeLink 1000 Plus (SR Research, Kanata, Canada) at a sampling rate of 1000 Hz. The remote mode was employed to capture data without requiring participants’ heads to be fixed on a chinrest, allowing for more natural, slight head movements. A 13-point calibration and validation procedures were conducted for each participant at the beginning of the experiment and checked before the start of each audio story. Only validations labeled as “good” by the EyeLink software were accepted. If a validation did not meet these standards, the procedure was aborted and re-run.

#### Preprocessing

Data analyses were performed in MATLAB (R2021b MathWorks). Pupil data was recorded and cleaned separately for each run. To assess how much the signals varied between the eyes, we calculated the Pearson’s correlation coefficient for each run. The average correlation across all runs and participants was strong indicating high similarity between the eyes’ recordings (r = 0.83, SE = 0.02). We linearly interpolated blinks and the other missing values due to non-detected pupil during the recording. We detected pupil values varying at a physiological unrealistically speed higher than 10 mm/s, flagged them as missing values and interpolated the data. Eye movements were regressed out of the pupil data, and the residual pupil signal was used for further processing. After trimming the pupil trace from the start to the end of the audio, the mean baseline period was subtracted from the data. Data was downsampled to 20Hz and a fourth-order Butterworth bandpass filter with cutoff frequencies of 0.1 Hz and 4 Hz was applied to the pupil signal.

### Text Preprocessing

Before passing the text through the language model, we performed preprocessing to ensure consistency in the input data. Punctuation marks were removed, and the text was converted to lowercase. Additionally, we replaced any instances of double hyphens or double spaces with single spaces to maintain proper spacing between words. These preprocessing steps were critical to reduce noise in the input data and ensure accurate lexical prediction calculations. The raw text from the podcast transcription, originally in RTF format, was converted to plain text using the striprtf package. Finally, extraneous whitespace was removed.

### Language modelling with GPT-2

We used GPT-2, a model from the transformer family^[44]^, specifically trained to compute the probability of each word given its preceding context. This model processes a sequence of tokens, either words or word-parts (e.g., sub-word tokens), into a sequence of conditional probabilities *P*(*w*_1_), *P*(*w*_2_∣*w*_1_), …, *P*(*w_T_*∣*w*_1_, …, *w*_*T* - 1_). The model’s architecture includes an embedding step that encodes each word as a vector (first hidden state ℎ_0_), which then serves as the input to a series of transformer blocks that sequentially update the hidden states ℎ_*t*_. At each block, a self-attention mechanism captures contextual dependencies between words. A (log-)softmax layer is applied to compute (log-)probabilities over target tokens.

The model can be summarized as:

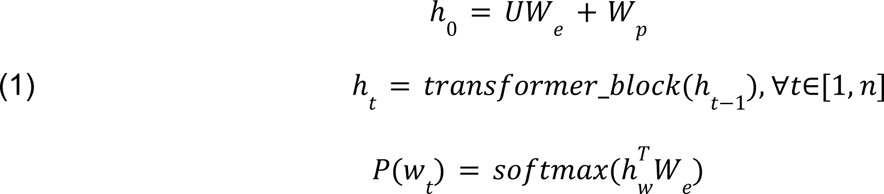

where *W_e_* is the token embedding matrix and *W_p_* is the positional embedding.

We used a smaller version of GPT-2 trained for the Dutch language (GroNLP/gpt2-small-dutch^[45]^). The model was loaded using the Huggingface transformers library and implemented in TensorFlow. The Dutch GPT-2 model consists of 345 million parameters. For each run, the preprocessed text of the podcast was tokenized using the Byte-Pair Encoding (BPE) scheme. Each tokenized segment was processed through a sliding window approach with a window size of 512 tokens. This allowed for efficient calculation of token probabilities while handling large chunks of text. During each pass through the window, the conditional probabilities of tokens given the preceding context were computed. Since GPT-2 handles fixed-length inputs (512 tokens), we implemented a sliding window approach to ensure that predictions spanned longer texts. Each 512-token window was processed sequentially, with overlapping windows to maintain context across segments.

### Base features

#### Speech envelope

The speech envelope for each stimulus was calculated using the mTRFenvelope function from the mTRF-Toolbox^[46]^ and then downsampled to 20 Hz to align with the pupil downsampled sampling rate. This speech envelope reflects the intensity of the speech signal over time, capturing energy fluctuations that correspond to phonemic and syllabic transitions. By incorporating the envelope feature, we effectively represent the acoustic response, allowing for a clear distinction between testing what aspect of pupil response is driven by pure acoustic processing and more higher-order linguistic processes (see next).

#### Word onset

The word onset regressor was created using a binary step function. This function takes a vector of word onset times and a time vector. For each onset, it identifies the closest index in the time vector and marks that position in a binary stimulus vector as 1. The resulting vector indicates the timing of each word onset. For the reversed speech condition, the same binary function was instead built from word offsets.

### Higher-order features

#### Word frequency

The word frequency regressor occurred at the same time as the word onset step function. To calculate word frequency, we used the Zipf-transformed word frequency value, calculated as Zipf = log₁₀((frequency + 1) / 44.106) + 3, where frequency is the raw count per million words from the SUBTLEX-NL corpus^[47]^. This logarithmic transformation standardizes the frequency scale, with values typically ranging from 1 to 7 (where 3 corresponds to approximately 44 occurrences per million), providing a more stable and interpretable measure than raw counts. Word frequency refers to a measure of familiarity, with more frequent words being more familiar and, therefore, easier to process than less frequent ones.

#### Semantic integration

Semantic integration difficulty was quantified for each word by comparing the embedding of that word as defined by the 320-dimensional GloVe word embeddings^[48]^ to the preceding context. Specifically, for each word w_n at position n in the transcript, a semantic context vector *c_n_* was constructed by averaging the embeddings of up to k = 10 preceding words (from indices max(1, n - window) to n - 1), and this context vector *c_n_* was compared to the target word’s embedding *v_wn_* using cosine similarity:

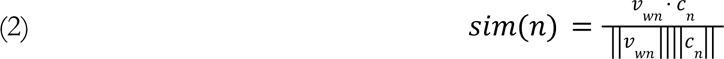

with a small constant (1e-10) added to the denominator to prevent division by zero. Integration difficulty was then defined as 1 - sim(n), yielding a regressor where higher values indicate greater difficulty integrating the current word into the evolving semantic representation. The first word of each story was assigned a similarity value of 1, as it had no preceding context to integrate with.

#### Surprisal and entropy

We quantified word-by-word unpredictability via lexical surprisal, calculated as the negative log-probability of a word given its preceding context:

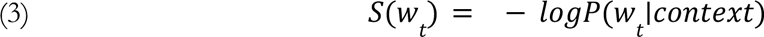

Where *logP*(*w_t_* ∣*context*) are the (log-)probabilities computed over target tokens in the softmax layer. The more probable a word is, based on its context, the less surprising.

In addition to surprisal, Entropy, shifted backward by one time step, was used to quantify the model’s uncertainty about the upcoming word:

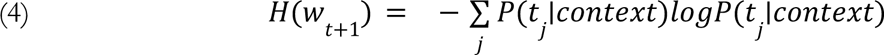

and the actual entropy value for word *w_t_* was obtained by applying the same one-step backward shift to the probability vector.

Entropy refers to the average level of surprise we anticipate for a word given its context. It is also known as expected surprise, as it reflects how surprising we expect a word to be. In contrast, surprisal measures how surprising a word actually is. While surprisal is a probabilistic measure of prediction error, entropy is a probabilistic measure of uncertainty. When entropy is high, predicting the next word is difficult; when it is low, the next word is much more predictable. Both the surprisal and entropy regressors were built from the word onset step function by substituting word onsets with the word surprisal or entropy.

For the reversed speech condition, all linguistic regressor vectors were flipped in order to match the reversed presentation order.

### TRF modelling

We used the Temporal Response Functions (TRFs) to model pupil data in response to acoustic and linguistic features^[46]^. The TRF approach operates within a multivariate linear regression framework, where lagged time series of input features serve as predictors. By modeling the response over time lags, TRFs help estimate the temporal profile of physiological responses to stimuli when there are no a priori hypotheses of their modulation by the regressors (such as the pupil response to word onset). This method enables the isolation of contributions to pupil modulation from distinct sources, including acoustic processing (e.g., speech envelope) and low/high-level lexical processing (e.g., word onsets/word frequency/surprisal/entropy),

Mathematically, the model can be expressed as:

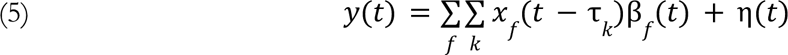

Here, *y*(*t*) represents the pupil response at time *t*, while *x_f_*(*t*) denotes the feature input at time *t* for feature *f*. The weights β_*f*_ (*t*) correspond to the TRF coefficients, capturing the temporal dynamics of how each feature influences pupil size (Figure 1). The term η(*t*) accounts for Gaussian noise in the data, representing variability not explained by the model.

To estimate the TRF coefficients, we employed ridge regression, which controls for overfitting by incorporating a regularization term. This approach is crucial given that features may exhibit autocorrelation, potentially leading to instability in coefficient estimation. The ridge regression formulation is given by:

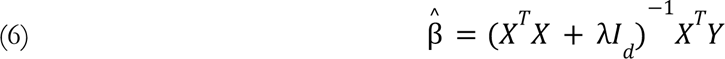

In this equation, *Y* is the matrix of measured pupil data, while *X* contains the lagged features’ timeseries. The regularization parameter λ ensures that the matrix *X^T^ X* remains well-conditioned, facilitating stable coefficient estimation. Given the slow response of the pupil, we chose time lags from -100 ms to 3000 ms.

To assess the effectiveness of our TRF model in reconstructing pupil data (model performance), we calculated the cross-validated Pearson correlation coefficient between the actual pupil responses and the reconstructed signals derived from the estimated TRFs (Figure 1). This correlation provides insight into how well our features account for the variance in pupil responses. Importantly, the TRFs were derived from a separate subset of the data (in a cross-validated manner), ensuring that model evaluation was not biased by overfitting to the training data.

To compare forward and backward story listening, we trained a separate linear encoder for each condition. For each participant, data from both the forward and backward versions of both stories were included. All models included an acoustic envelope regressor to capture low-level acoustic effects and a word-onset stick function (baseline model). A single higher-level lexical feature was included as a parametric modulation of the word-onset regressor, ensuring that all models contained the same number of regressors and differed only in the highest-level feature: (1) envelope, word onset, and word frequency; (2) envelope, word onset, and surprisal; (3) envelope, word onset, and entropy; (4) envelope, word onset, and semantic integration. For each model, we compared temporal response functions (TRFs) and model performance between the forward and backward conditions. Both pupil data and all regressors except the word onset regressor were z-scored before entering the model.

### Statistical analyses

#### TRF analysis

To statistically compare temporal response functions (TRFs) between forward and backward conditions, we employed a threshold-free cluster enhancement (TFCE) approach combined with permutation testing^[49]^. Given the paired-samples design (both conditions measured within each participant), we computed the participant-wise difference between conditions (forward − backward) and derived a one-sample t-statistic at each time point against zero. To construct the null hypothesis distribution assuming no difference between conditions, we applied a sign-flipping procedure: for each permutation (5,000 iterations), we randomly multiplied each participant’s difference waveform by either +1 or −1 (i.e., permuting the sign of the condition difference within participant), recomputed the t-statistic, and calculated the TFCE scores separately for positive and negative t-value clusters. The maximum TFCE score across time points and tails per permutation served as the null distribution. For the observed data, we computed the same TFCE scores and obtained nonparametric p-values by comparing the observed TFCE at each time point against this null distribution, controlling the family-wise error rate (FWER) across time. TFCE parameters were set to E = 1, H = 2, with a step size (dh) of 0.1 and a minimum threshold of 0^[50]^.

#### Model performance

Prediction performance as measured by cross-validated Pearson correlation coefficients, i.e., reconstruction accuracy, of forward and reverse TRF models was compared using paired two-tailed t-tests across participants for each model, for example each channel or time-lag. To correct for multiple comparisons, the false discovery rate (FDR; Benjamini–Hochberg) was applied with α = 0.05.

## RESULTS

We recorded eye-movements while participants listened to podcasts. Each podcast contained two stories, with both stories played twice. Additionally, one of the two stories was also reversed (i.e. resulting in reversed, nonsense speech), and was also played twice. This resulted in six runs of audio per participant - two stories with normal (forward) and one with reverse speech (Figure 1). The order of three stories was randomized. All three stories were presented once before repeating them (thus order for runs 4-6 was the same as for runs 1-3).. Pupil signal was recorded from both eyes and then averaged. Eye movements were regressed out, and the resulting residual pupil signal was used for further analysis.

Our primary prediction was that the pupil responds to contextual lexical surprisal in stories. We quantified contextual surprisal from a generative language model (Figure 1, Dutch GPT^[45]^). We additionally examined lower-level, non-contextual word features (e.g., word frequency) and semantic integration difficulty quantified via sliding-window cosine similarity over static GloVe embeddings, which capture word co-occurrence statistics. Unlike lexical surprisal, which captures moment-by-moment violations of sequential expectations, semantic integration reflects a cumulative, context-insensitive fit to the preceding semantic field. Entropy (expected surprise) was also quantified using GPT-2. For the reverse condition, all linguistic regressors were time-reversed, and word onsets were defined using the word offsets from the corresponding forward-condition stimuli. To assess how these features reflect pupil dilation, we employed a temporal response function (TRF) modeling approach, examining estimated TRF weights over time and reconstruction accuracies via cross-validated Pearson’s correlation (see Methods for details).

### Linguistic features successfully predict pupil responses to forward speech

To investigate model performance, we calculated cross-validated Pearson correlations between actual and predicted pupil responses (Figure 1). Coefficients were estimated from a separate data subset to avoid overfitting. We compared five models: one baseline model with the low-level features and four models combining the baseline model with one higher-level feature: (1) word frequency, (2) surprisal, (3) entropy, or (4) semantic integration. For each model, we compared performance between the forward and backward conditions.

Across all five models, reconstruction accuracies were consistently higher for forward compared to backward listening (Figure 2). This forward advantage reached significance in four of the five models: the baseline model (envelope +word onset; forward > backward: t(42) = 2.40, p = .026), the envelope + word onset + word frequency model (forward > backward: t(42) = 2.76, p = .017), the envelope + word onset + surprisal model (t(42) = 3.61, p = .004), and the envelope + word onset + semantic integration model (t(42) = 2.68, p = .017). The strongest effect was observed for the model including lexical surprisal, which yielded the most robust forward–backward difference. In contrast, the envelope + word onset + entropy model showed a marginally significant difference between conditions (t(42) = 1.84, p = .073). Direct comparisons between forward models did not reveal significant differences in prediction accuracy, suggesting that while the surprisal confers a particularly strong forward encoding advantage, the overall pattern reflects a general enhancement of model performance for forward linguistic structure.

**Figure 2.**
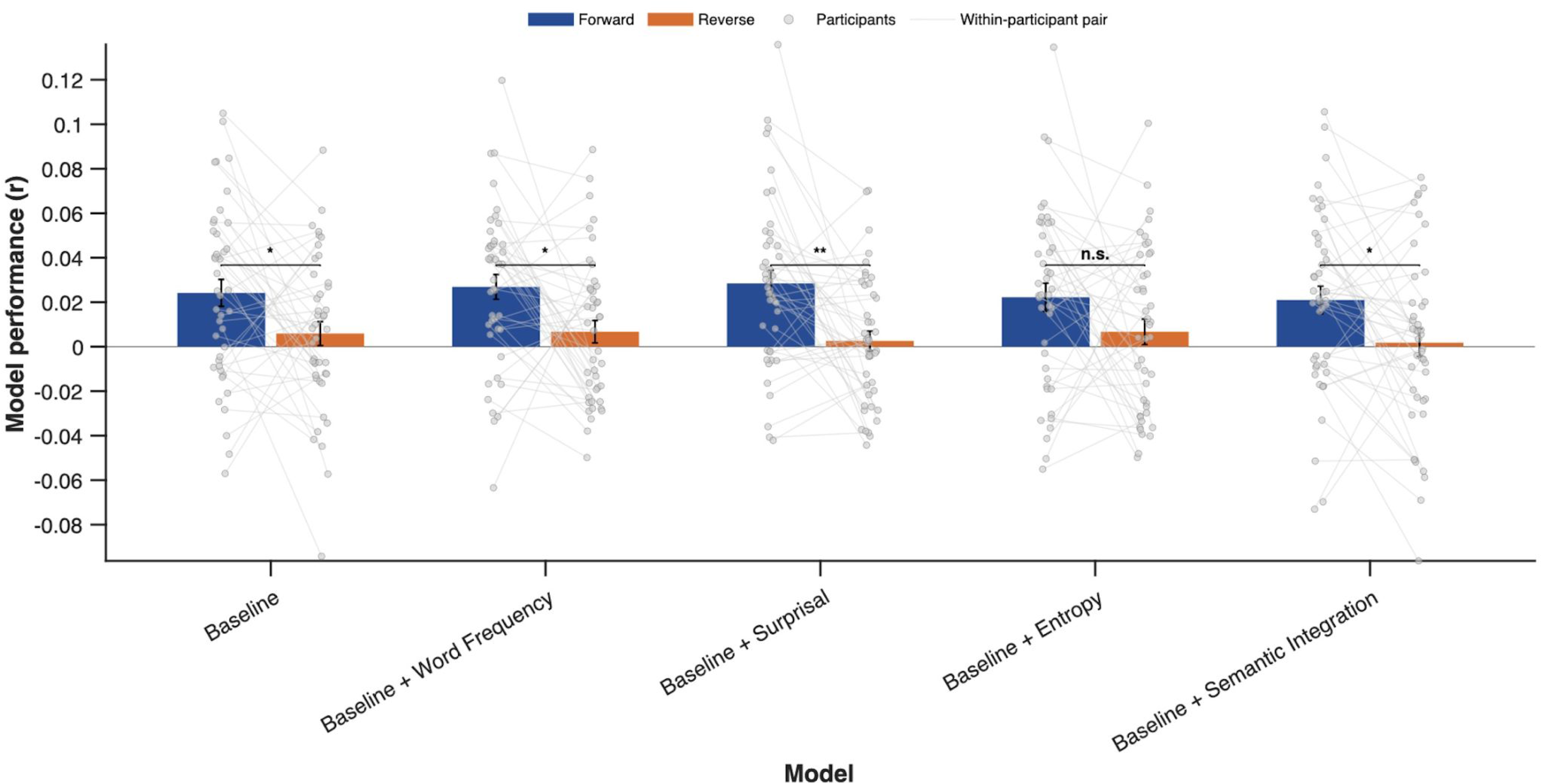
Model performance for forward and reverse story listening. Models performance was consistently higher for forward than backward listening across all five models. Forward models outperformed backward models for four of the five linguistic predictors, with the most robust forward advantage observed for the surprisal model. The entropy model also showed a marginal forward tendency, though its effect was weaker than the others. Direct comparisons between forward models revealed no significant differences in prediction accuracy. *p < .05, **p < .01

### Pupil responds to lexical surprisal during naturalistic listening

To characterize the temporal profile of how acoustic and linguistic features influence pupil dilation, we examined temporal response functions (TRFs) across all five models. A TRF can be understood as a linear filter that describes, for a given predictor, how a unit change in that feature modulates the pupil response across a continuous range of time lags. Because the envelope + word onset + surprisal model performed numerically best in the preceding comparison, its TRFs are presented and serve as the primary focus; TRFs for the remaining models are reported in the supplementary results section. For each model, TRFs for the envelope, word-onset, and lexical regressors were extracted and compared between forward and backward listening conditions. Statistical significance was assessed using threshold-free cluster enhancement (TFCE) with sign-flipping permutation tests (5,000 iterations). The null distribution was constructed by randomly multiplying each participant’s condition difference waveform (forward − backward) by either +1 or −1 per permutation (i.e. flipping the sign of the condition contrast), recomputing the t-statistic, and calculating TFCE scores separately for positive and negative t-value clusters. The maximum TFCE score across time points and tails per permutation served as the null distribution, and observed TFCE scores were compared against this distribution to obtain nonparametric p-values controlling the family-wise error rate (FWER) across time. For each significant cluster, the reported p-value corresponds to the maximum absolute t-value within that cluster (i.e., the cluster peak).

### Lexical surprisal

Consistent with our primary prediction, the TRF for the lexical surprisal regressor revealed a significant difference between forward and backward conditions within a time window of 1700–2350 ms following word onset (p = 0.0082) (Figure 3). In this interval, the pupil response showed greater dilation for words with higher contextual surprisal, a pattern that corresponds to the surprisal-evoked dilation effect previously reported in the literature. The effect emerged within a late time window (1700–2350 ms), which is consistent with the temporal profile of surprisal-related pupil responses documented in previous work. Overall, this pattern is consistent with the view that pupil diameter during naturalistic listening tracks context-dependent prediction error rather than word familiarity or uncertainty.

**Figure 3.**
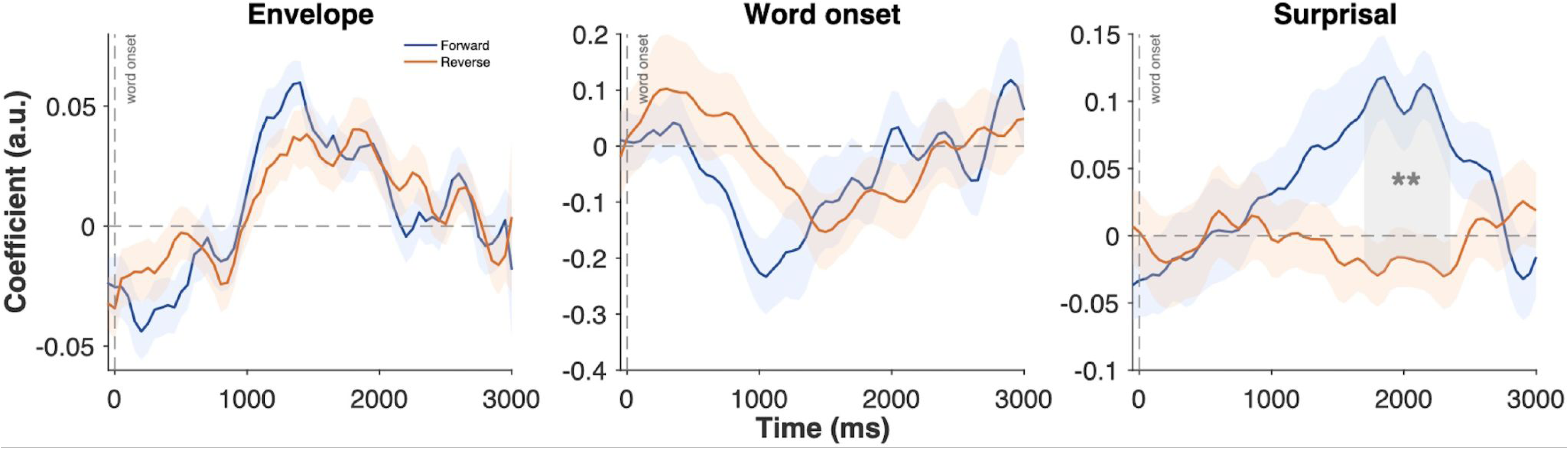
Temporal response functions (TRFs) for the surprisal model. Left: envelope TRF. Center: word-onset TRF. Right: lexical surprisal TRF. For the lexical surprisal regressor, the forward condition elicited robust pupil dilation responses compared to the backward condition, with significant differences localised to a time window of 1700–2350 ms post-word onset, indicating a pupil response to contextual surprisal (unexpectedness). Grey-shaded areas denote time windows of significant effects identified via cluster-based permutation testing with TFCE (p < .01).

## DISCUSSION

The present study investigated whether the pupil responds to surprisal in a completely nonvisual setting, during listening to an auditory narrative. We further investigated the nature of this surprisal, whether it tracks word frequency or semantic integration, thus responding to local context, or more broadly tracks the context of a narrative. This was done in a continuous naturalistic way as participants listened to an ongoing narrative, as opposed to more common trialwise examinations of pupil responses^[51]^. To this aim, we employed TRF modeling^[46]^ to investigate the dynamics of pupil responses during continuous speech. Predictors were derived from the text of the podcast stories. We compared one baseline model (i.e. acoustic envelope and word onset) and four matched-complexity models (i.e. baseline model and one higher-order linguistic feature parametrically modulating word onset: word frequency, lexical surprisal, entropy, or semantic integration) applied to pupil data recorded while participants listened to forward and backward versions of the same stories. Our primary finding is that lexical surprisal significantly modulates pupil diameter during a late time window (1700–2350 ms) in forward speech. The cluster-based comparison revealed a positive cluster reflecting a surprisal effect that was observed during forward speech but not during backward speech, confirming that this effect is specifically driven by the processing of meaningful linguistic structure, rather than by low-level acoustic properties of the speech signal. In contrast, we observed no reliable modulation for word frequency, entropy, or semantic integration. This surprisal-evoked dilation effect is consistent with our prediction that the pupil responds to contextual prediction error during naturalistic language comprehension.

Why would the pupil selectively track contextual surprisal during naturalistic listening? The answer likely lies in the functional properties of the locus coeruleus-norepinephrine (LC-NE) system, which is closely coupled to pupil dilation and plays a fundamental role in regulating attention and learning in response to environmental change. LC-NE neurons increase their firing rate phasically when unexpected events occur, and this response habituates with repetition, a pattern that closely resembles the signature of a prediction error signal^[52]^. Pupil dilation has been consistently shown to reflect this phasic LC-NE activity, making it a reliable peripheral marker of surprisal-related arousal^[9,17,18,22-28]^.

The temporal profile of this effect is also particularly informative. The late window is consistent with the well-established sluggishness of the pupillary response. In humans, the pupil response begins around 1000 ms after stimulus onset^[40,53]^ and peaks at latencies between 1500 and 2500 ms comparable to those reported in prior work investigating surprisal in both tone listening^[18]^ and continuous speech^[33]^. Indeed, cortical markers of linguistic surprisal have been observed around 600 ms post-stimulus^[30,38]^; once the roughly one-second neuromodulatory-to-pupil lag is taken into account, these neural effects likely map onto the late dilation window we observe.

Recent work by Gehmacher et al.^[33]^ has shown that during story listening, cortical and pupil-linked systems track contextual predictions in distinct but complementary ways. The auditory cortex maintained a continuous, graded map of lexical surprisal, while tonic pupil diameter and brainstem-localised responses additionally showed a preference for a rectified linear (ReLU) formulation of semantic prediction error, responding selectively when meaning-relevant errors exceeded a recent contextual baseline, which the authors linked to model updating and attentional reorientation. Importantly, this gating signature was specific to semantic prediction error: for lexical surprise, pupil responses were captured equally well by the graded formulation, with no significant ReLU advantage. Our findings converge with this graded component of their results. We find that the pupil responds to lexical surprisal across a broad, continuous range of magnitudes, responding to all levels of surprisal rather than only to the most surprising events, consistent with Gehmacher et al.’s^[33]^ observation that pupil-linked arousal tracks lexical surprise in a graded fashion.

This framework fits with our finding that the pupil responds to surprisal but not to word frequency or static semantic similarity. Word frequency is a static property of the lexicon; encountering a low-frequency word does not in itself constitute a violation of an ongoing prediction, and therefore may not trigger the phasic LC-NE response that drives pupil dilation. This finding underscores a fundamental shift in processing dynamics when moving from isolated words to continuous, meaningful speech. In single-word paradigms, the cognitive system must rely primarily on lexical access mechanisms, making word frequency (a direct index of lexical difficulty) a potent driver of pupil dilation^[40,41]^. Low-frequency words are encountered less often, possess higher resting activation thresholds, and require greater cognitive effort to recognize and retrieve from the mental lexicon; this increased lexical access difficulty is what the pupil has been shown to track in isolated word tasks. However, in a rich narrative context, the preceding sentence and discourse create a powerful semantic frame that pre-activates features of upcoming words, reducing or even eliminating the need for a purely frequency-based lexical search. Critically, under such strongly constraining contexts, surprisal subsumes frequency effects: a low-frequency word can be highly predictable when it fits the contextual model, while a high-frequency word can be highly surprising if it violates expectations. The brain’s processing effort, indexed by the pupil, is therefore driven by this contextual fit rather than the word’s prior probability in isolation (frequency). This interpretation aligns with recent evidence that the pupil’s sensitivity to listening effort is markedly attenuated during naturalistic story listening compared to isolated sentence paradigms^[32,54]^.

Similarly, semantic integration captures broad meaning relationships that lack the temporal precision and contextual sensitivity needed to engage the brainstem-pupil axis. Our results revealed that although the semantic integration regressor closely resembled surprisal in shape, it did not significantly distinguish forward from backward speech. This likely reflects a limitation of GloVe embeddings, which are static, non-contextual representations derived from global corpus co-occurrence statistics: they capture broad semantic relatedness but lack sensitivity to the dynamic, moment-by-moment shifts in meaning that arise from unfolding linguistic context. In contrast, surprisal from GPT-2 is inherently contextual, quantifying the processing cost of a word given its specific preceding discourse. Consistent with this distinction, Michaelov et al.^[37]^ showed that surprisal from a contextual language model (GPT-3) significantly outperformed GloVe-based similarity in predicting N400 amplitudes, suggesting that the brain’s response to language is better characterized by dynamic predictability than by static semantic fit.

Because surprisal reflects the fit between a word and its specific prior context, it signals a genuine deviation from the listener’s active model of the narrative, the kind of event that calls for model updating, reorienting attention, and facilitating learning^[55]^. The pupil, therefore, appears to track the cognitive cost of updating a predictive model during naturalistic listening. This suggests that in passive listening, the pupil is not a monolithic index of lexical difficulty, nor is it broadly sensitive to static semantic relatedness. Rather, it is selectively attuned to the dynamic interplay between contextual prediction and event-level narrative structure, a process for which surprisal serves as the more informative metric.

The absence of a reliable entropy effect can be understood within the broader context of how the pupil-linked arousal system responds to different forms of uncertainty. Entropy, as operationalized here, quantifies the model’s expected surprisal, the average level of surprise anticipated for a word given its preceding context, rather than the actual surprisal elicited when the word is encountered. While previous work has demonstrated pupil sensitivity to uncertainty during decision making and auditory sequence learning, these effects have typically emerged under conditions involving environmental volatility or contextual shifts that necessitate model updating^[18,56]^. In the present task, by contrast, the statistical structure of the language and the narrative itself were not subject to sudden, experimenter-manipulated changes that would render the listener’s predictive model invalid. Under these conditions, entropy reflects the uncertainty about upcoming linguistic input before the word arrives, that is, how difficult it is to predict the next word given the context. This expected surprise, however, does not itself constitute a violation of an ongoing prediction; it describes the uncertainty that exists prior to the event and it may influence the overall computational load on the language system, but it does not necessarily trigger the phasic arousal response that pupil dilation indexes. This dissociation is consistent with the proposal that pupil-linked arousal reflects the cognitive cost of updating a predictive model in response to violations, rather than the uncertainty that precedes those violations^[24]^.

Our findings position the pupil as a useful marker of prediction error in naturalistic language comprehension. Whereas previous pupillometric work established sensitivity to auditory regularity violations in tightly controlled paradigms^[17,18]^, the present results show that this sensitivity extends to lexical-level prediction error during continuous, ecologically valid speech. This broadens the scope of pupillometry as a tool for investigating predictive processing outside the laboratory, and provides a low-cost, non-invasive peripheral measure with which hierarchical predictive coding accounts can be tested^[20,21,24]^.

However, several limitations should be considered. TRF modeling approach, while powerful for capturing the temporal dynamics of pupil responses, assumes linear relationships between features and the pupil signal, which may not fully capture the nonlinear nature of cognitive processing. Second, while we controlled for low-level acoustic features through the backward speech condition, other potential confounds (i.e. prosodic or rhythmic structure) may have influenced the forward–backward differences observed. Future work should examine different components of predictions, such as syntactic categories and phonemes, to determine whether the pupil also responds to these features or is more specifically related to the semantic content of words. Syntactic and phonemic predictions modulate early, focal temporal responses, whereas semantic predictions engage later, distributed activity^[30,38]^. Investigating these hierarchical levels of prediction within the pupillary system could provide further insight into the role of the LC-NE system in language comprehension. Furthermore, our experimental design, in which participants listened to the same story twice, introduces a potential confound of stimulus familiarity or learning effects. It is possible that predictive processes differed between the first and second presentations, with repeated exposure allowing listeners to form more precise expectations about upcoming linguistic content. Unfortunately, our current dataset lacked sufficient statistical power to systematically investigate these temporal dynamics across sessions, and future studies should explicitly manipulate exposure (e.g., by comparing single versus repeated presentations) to disentangle learned anticipation from online predictive processing. Relatedly, examining individual differences in working memory capacity, language proficiency, or learning rate could reveal how listeners adaptively allocate resources under varying levels of contextual constraint, and whether these cognitive factors modulate the forward–backward pupil differences we observed.

A key strength of our paradigm, and a promising direction for future work, lies in its fully non-visual, free-gaze design. By having participants listen in darkness without fixation instructions, we avoided well-known visual confounds inherent to pupillometry, including luminance fluctuations, contrast adaptation, and accommodative effort, while preserving the ecological validity of connected speech comprehension. This approach may be especially valuable for populations in which sustained fixation or task compliance is difficult, as it enables passive, unconstrained assessment of prediction-related cognitive effort. Future studies could extend this framework to developmental populations, such as infants or young children, as well as clinical cohorts, to investigate whether predictive processing during continuous speech comprehension can be reliably indexed via the pupil response in the absence of visual demands.

## Acknowledgements

C.T. was funded by European Research Council (ERC-AdG 832813-GenPercept awarded to David Charles Burr). This publication is part of the project “Do we use our eyes to think?” with file number VI.Veni.231G.048 of the research programme NWO Talent Programme Veni which is (partly) financed by the Dutch Research Council (NWO) awarded to E.B.

---

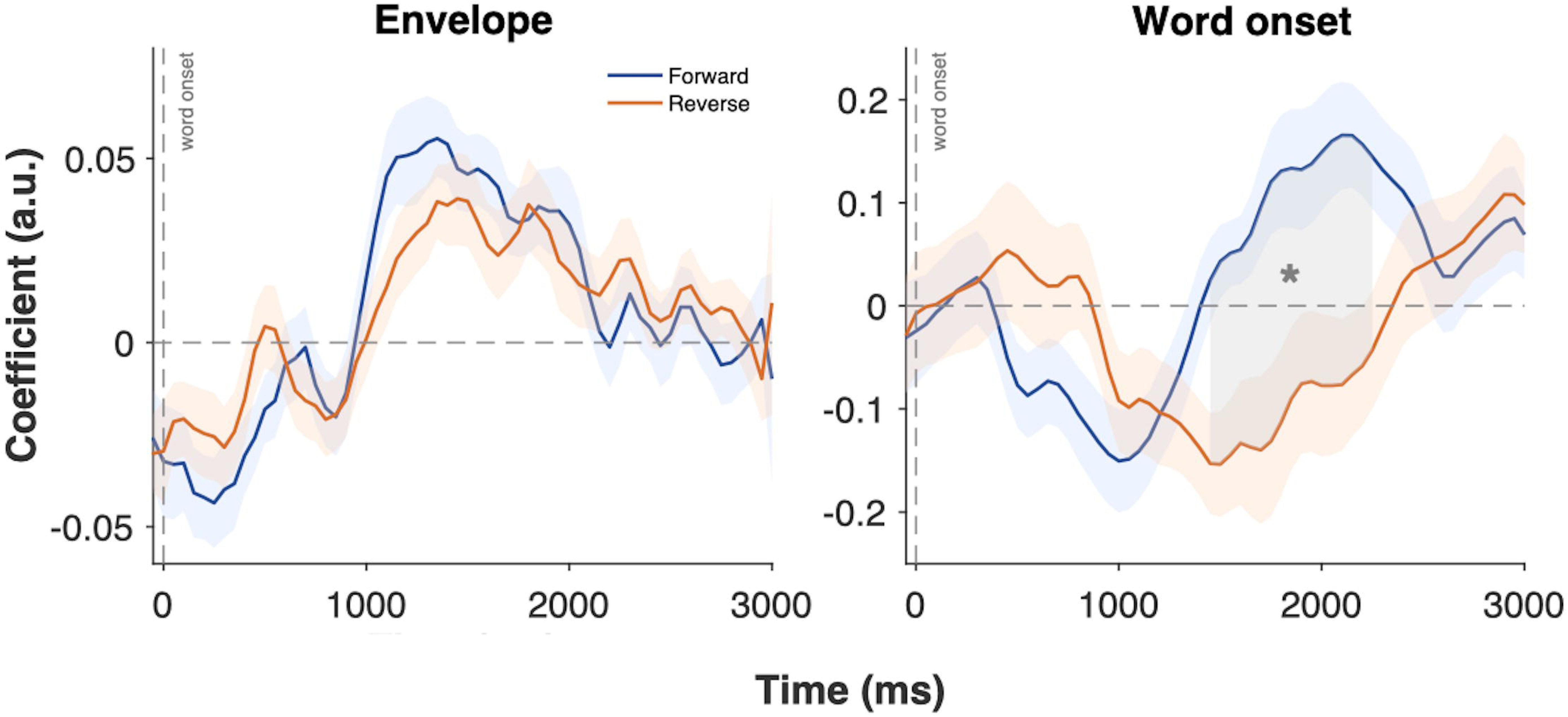

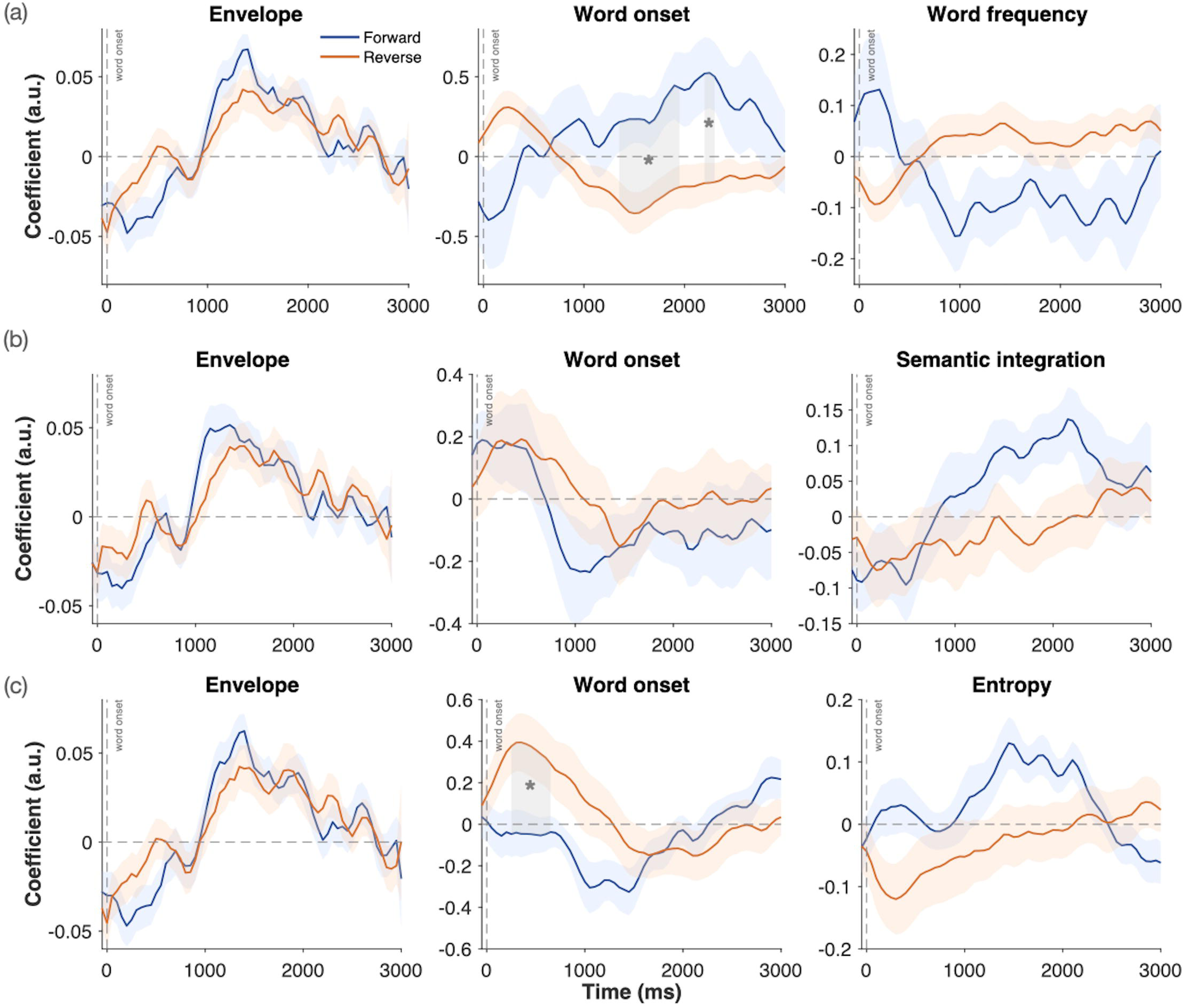

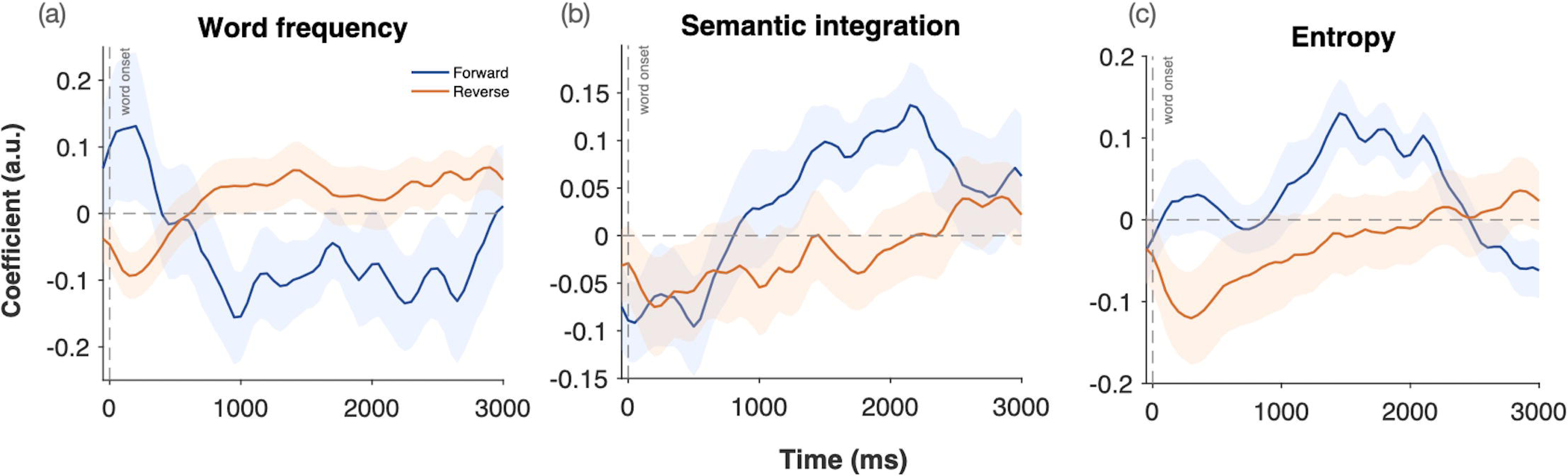

