## Supplementary material for "Lexical surprisal drives pupil dilation during continuous story listening"


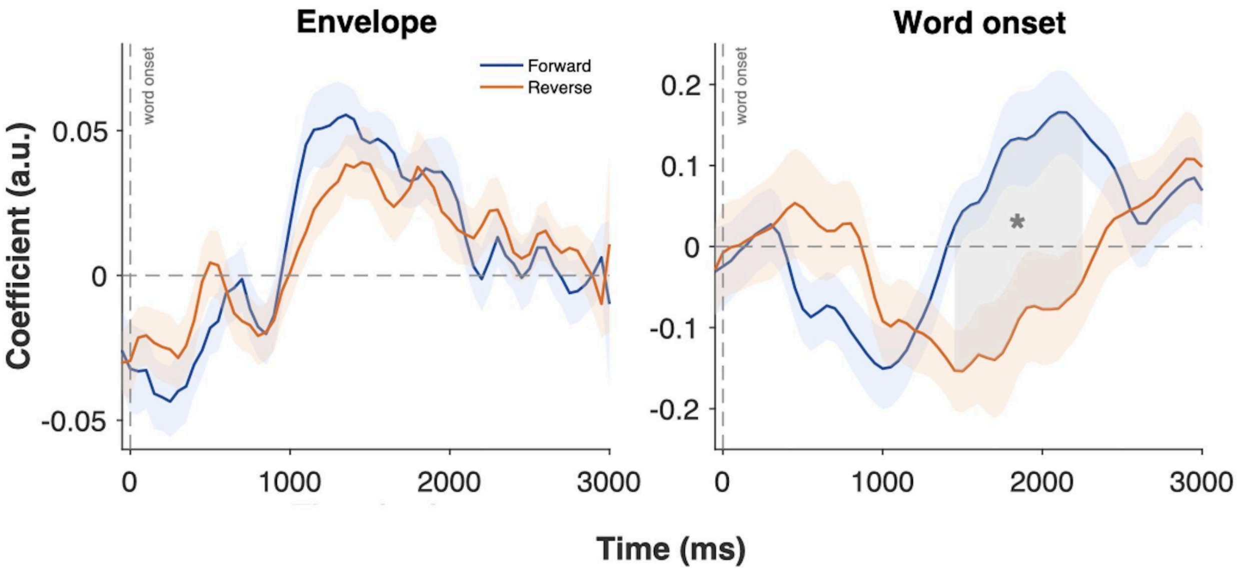


**Figure 1. TRFs for the baseline model.** Left: envelope TRF. Right: word-onset TRF. For the word onset regressor, the forward condition elicited robust pupil dilation responses compared to the backward condition, with significant differences localised to a time window of 1450-2250 ms post-word onset. Grey-shaded areas denote time windows of significant effects identified via cluster-based permutation testing with TFCE ($p < .01$).

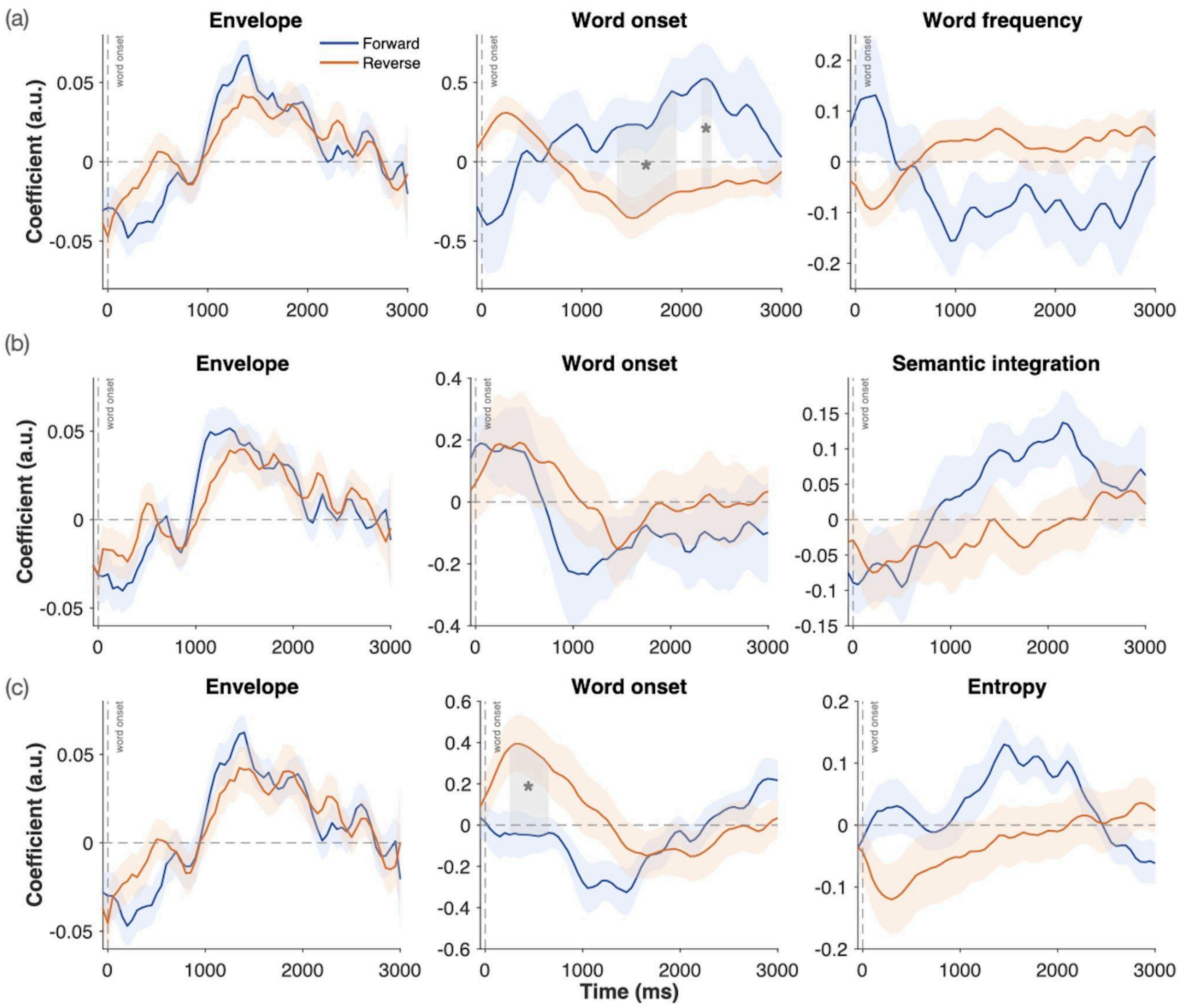


**Figure 2. TRFs for all regressors across three models**. (a) Envelope + word onset + word frequency model: significant forward–backward differences for the word-onset regressor were observed from 1350–1950 ms ($p = 0.0156$) and 2200–2300 ms ($p = 0.0478$), with forward speech eliciting more positive beta coefficients than backward speech within these windows. (b) Envelope + word onset + entropy model: significant forward–backward differences for the word-onset regressor emerged from 250–650 ms ($p = 0.0348$), with backward speech eliciting more positive beta coefficients than forward speech. (c) Envelope + word onset + semantic integration model: no significant differences were observed for any regressor. No significant forward–backward differences were found for the word frequency, semantic integration, or entropy regressors in any time window.

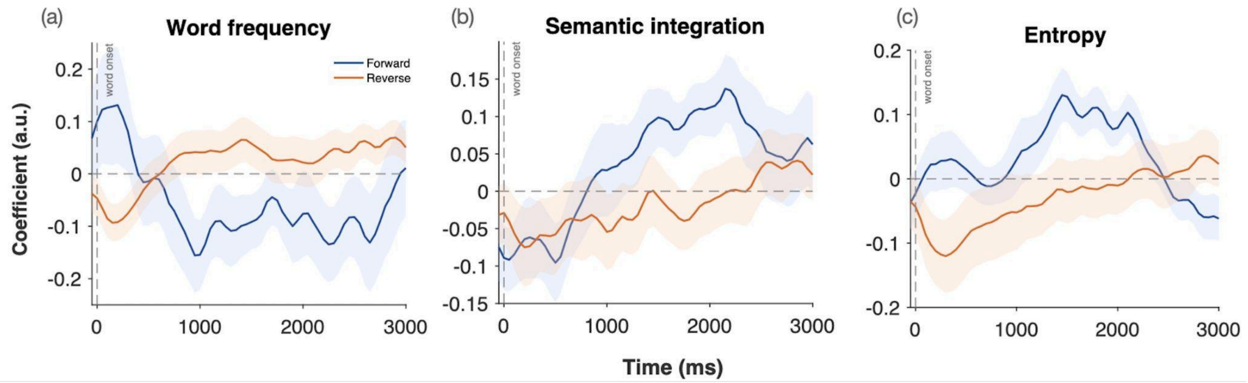


**Figure 3. TRFs for linguistic regressors across three models**. (a) Word frequency; (b) Semantic integration; (c) Entropy. No significant forward–backward differences were observed for any of the linguistic regressors.

### Baseline model

The base model comprises envelope and word onset predictors. As expected, the envelope predictor did not differ between forward and backward conditions, confirming that the low-level acoustic properties of the speech signal are preserved in the backward condition. On the other hand, we found a strong dilation response to word onset only when participants were listening to forward speech the 1450-2250 ms time window ($p = 0.0192$), indicating that the pupil dilation response to the onset of a word depends on whether the speech is intelligible.

### Word onset effects in the linguistic models

All linguistic regressors were constructed from the word onset stick function and are therefore correlated with the word onset regressor. Differences between models in the word onset effect may thus reflect how much of this shared onset-locked variance each predictor accounts for, which may depend on the predictor's strength and on the similarity of its response shape to the word onset response.

The word onset effect was absent in both the surprisal model and the semantic integration model (Figures 1d, 1f), in which the semantic similarity regressor itself was not significant (Figure 1e). In both models, the linguistic TRF was similar in shape to the word onset response. For surprisal, this may reflect the convergence of two factors: it is the strongest linguistic predictor of the pupil response, and its TRF resembles the onset response, so it may account for much of the variance shared with word onset. The semantic integration model suggests that shape similarity alone may be sufficient to obscure the onset effect: when two correlated regressors have similarly shaped responses, the condition difference in their shared time window may be divided between them, increasing the variance of both coefficient estimates and potentially masking effects. The entropy TRF was also similar in shape to those of surprisal and semantic integration (Figure 1g), and no late word onset window differed between conditions. A significant early window did emerge, however, from 250 ms to 650 ms ($p = 0.0360$) (Figure 1b), with larger coefficients for backward than forward speech. This latency falls within the range of the early, phasic component of the pupil response, which may index a stimulus-driven orienting response rather than the slower dilation associated with cognitive effort[57,58].

In the envelope + word + word frequency model, the word onset regressor resembled the baseline pattern, with significant windows from 1350 ms to 1950 ms ($p = 0.0176$) and from 2200 ms to 2300 ms ($p = 0.0472$) (Figure 1a); forward speech elicited more positive coefficients than backward speech, at latencies consistent with typical pupil dilation responses. The word frequency TRF differed from the other linguistic TRFs in shape, showing a marked negative inflection rather than a dilation-like positive waveform. This distinct shape may explain why the word onset effect remained significant: the frequency regressor may not compete for onset-locked variance, leaving the word onset response largely unaffected. The negative polarity is consistent with smaller pupil dilations for more frequent words[40,41].

### Word frequency, semantic integration, and entropy

No significant differences between forward and backward conditions were observed for the word frequency regressor in any time window (Figure 4a). Similarly, the entropy (Figure 4b) and the semantic integration (Figure 4c) regressors did not yield any significant TFCE clusters.
